# The transcriptome of playfulness is sex-biased in the juvenile rat medial amygdala: a role for inhibitory neurons

**DOI:** 10.1101/2024.09.11.612456

**Authors:** Ashley E. Marquardt, Mahashweta Basu, Jonathan W. VanRyzin, Seth A. Ament, Margaret M. McCarthy

## Abstract

Social play is a dynamic behavior known to be sexually differentiated; in most species, males play more than females, a sex difference driven in large part by the medial amygdala (MeA). Despite the well-conserved nature of this sex difference and the importance of social play for appropriate maturation of brain and behavior, the full mechanism establishing the sex bias in play is unknown. Here, we explore “the transcriptome of playfulness” in the juvenile rat MeA, assessing differences in gene expression between high- and low-playing animals of both sexes via bulk RNA-sequencing. Using weighted gene co-expression network analysis (WGCNA) to identify gene modules combined with analysis of differentially expressed genes (DEGs), we demonstrate that the transcriptomic profile in the juvenile rat MeA associated with playfulness is largely distinct in males compared to females. Of the 13 play-associated WGCNA networks identified, only two were associated with play in both sexes, and very few DEGs associated with playfulness were shared between males and females. Data from our parallel single-cell RNA-sequencing experiments using amygdala samples from newborn male and female rats suggests that inhibitory neurons drive this sex difference, as the majority of sex-biased DEGs in the neonatal amygdala are enriched within this population. Supporting this notion, we demonstrate that inhibitory neurons comprise the majority of play-active cells in the juvenile MeA, with males having a greater number of play-active cells than females, of which a larger proportion are GABAergic. Through integrative bioinformatic analyses, we further explore the expression, function, and cell-type specificity of key play-associated modules and the regulator “hub genes” predicted to drive them, providing valuable insight into the sex-biased mechanisms underlying this fundamental social behavior.

## Introduction

Social play, also known as rough-and-tumble play or play-fighting, is a complex, rewarding behavior expressed by juveniles across most mammalian species. While its exact function is debated, converging evidence suggests play serves a vital role in appropriate brain development, as animals socially isolated as juveniles exhibit various behavioral impairments in adulthood, including impaired social behavior (Hol et al., 1999; Van den Berg et al., 1999; Von Frijtag et al., 2002; Marquardt et al., 2023), dysregulated cognition (Einon et al., 1978, Baarendse et al., 2013; Yusufishaq & Rosenkranz, 2013) and increased anxiety- and depression-like behavior (Parker, 1986; Wright et al., 1991; Arakawa et al., 2003; Leussis & Anderson, 2008; Lukkes et al., 2009; Cuesta et al., 2020).

Importantly, a core feature of social play is its sex bias. From rats to cats to humans, strong sex differences in play are observed in nearly all species that exhibit this behavior, whereby males play more frequently and more intensely than females (reviewed in VanRyzin et al., 2020a). This sex difference appears to be strongly driven by the medial amygdala (MeA), as MeA lesions do not abolish play but instead remove the sex difference by specifically lowering male play levels to that seen in females (Meaney et al., 1981), and testosterone implants to the neonatal MeA increase the play of female juveniles to that seen in males (Meaney & McEwen, 1986). Accordingly, play deficits are core features of various neurodevelopmental disorders such as autism spectrum disorder and attention deficit/hyperactivity disorder (Alessandri, 1992; Jordan, 2003; Mahendiran et al., 2019), many of which also exhibit a sex bias in prevalence and severity (Polyak et al., 2015). By investigating the causes and consequences of sex differences in play, we gain insight as to how brain sex differences are typically established in a manner that may prove useful for understanding why boys and girls are differentially at risk for these disorders.

While there is a reliable sex difference in average play frequency and intensity in many mammalian species, it is essential to note that there is a high degree of individual variability in playfulness, even in inbred laboratory animals, not unlike that seen for many other behaviors (Tuttle et al., 2018; Rudolová et al., 2022). Although males play more frequently *on average*, some females play at the level of even high-playing males, while some males play at the level of low-playing females. These individual differences largely persist over time; play levels in rats, analyzed both at the level of the pair and the level of the individual, have been found to be remarkably consistent throughout adolescence and across motivational contexts (Pellis & McKenna, 1992; Argue & McCarthy, 2015; Lampe et al., 2017). Here, we exploit the power of these individual differences in playfulness to determine whether social play is associated with the same or different gene expression profiles in the MeA of male and female rats. We find that the latter is true: the gene signatures in the MeA associated with playfulness are largely distinct in juvenile males and females. We combine these analyses with parallel single-cell RNA-sequencing analyses of the neonatal amygdala to better understand how this transcriptional sex difference is established, discovering that inhibitory neurons are a major locus of sex-biased gene expression in this region. Finally, we explore the predicted cell-type specificity, function, and key hub genes within representative play-associated modules of interest using bioinformatics, phenotyping play-active cells and those expressing module hub genes in the juvenile MeA using RNAscope®. To our knowledge, this is the first study assessing gene expression patterns associated with play in both sexes, aiding in our understanding of the genesis and relevance of sex differences in this essential adolescent behavior.

## Materials and Methods

### Experimental subjects

Adult Sprague-Dawley rats (Charles River Laboratories, Wilmington, MA) were maintained on a 12:12h reverse light-dark cycle with *ad libitum* access to food and water. Animals were mated in our facility and allowed to deliver normally under standard laboratory conditions, with the day of birth designated as postnatal day 0 (P0). Animals of both sexes, balanced across multiple litters, were used in this study. All animal procedures were performed in accordance with the regulations of the Institutional Animal Care and Use Committee at the University of Maryland School of Medicine.

### Social play behavioral testing

Animals were weaned on P21 and housed in same-sex sibling pairs. Starting on P26, social play behavior was assessed as described previously (VanRyzin et al., 2020b). Same-sex non-sibling pairs of animals were placed in an enclosure (49 x 37 cm, 24 cm high) with TEK-Fresh cellulose bedding (Envigo, Indianapolis, IN). Animals were allowed to acclimate to the arena for 2 min, after which their interactions were video recorded for 10 min before being returned to their home cages. This procedure was repeated once daily from P27-29, using the same pairs of animals each day. All behavioral testing was performed during the dark phase of the light-dark cycle under red light illumination. Videos were scored offline to quantify the number of pounces, pins, and boxing behaviors, summed together as the total play behaviors exhibited in the test. Animals that exhibited an average play score that was in the top third or bottom third per sex were included in later RNA-seq analysis, resulting in four groups: low-playing females (FLO), high-playing females (FHI), low-playing males (MLO), and high-playing males (MHI).

### Bulk RNA-sequencing (RNA-seq)

On P30 (∼24 hours after the final play bout), juveniles (n=6 per group) were deeply anesthetized with Fatal Plus (Vortech Pharmaceuticals, Dearborn, MI) and decapitated. Brains were removed and placed on ice, and the amygdala of both hemispheres was dissected out and immediately flash frozen. Tissue was stored at −80°C until it was prepared for RNA extraction. RNA was extracted using the RNeasy Kit (Qiagen, Hilden, Germany), following the manufacturer’s instructions, including column treatment with DNase I (Qiagen) to remove contaminating DNA. Bulk RNA-seq was performed by staff of Maryland Genomics at the Institute for Genome Sciences of the University of Maryland School of Medicine. Libraries were prepared using a NEBNext Ultra Directional RNA prep kit (New England Biolabs, Ipswich, MA) using the manufacturer’s protocol. Samples were sequenced across 4 channels using the 75 base pair paired-end protocol on an Illumina HiSeq 4000 sequencer, targeting 50 million read pairs per sample.

### Bulk RNA-seq analysis

Sequencing reads were aligned to the *Rattus norvegicus* genome (Rnor v7.2) using HISAT2 (Kim et al., 2015), and tables of read counts per transcript were generated with HTSeq (Anders et al., 2015). Multi-dimensional scaling and hierarchical clustering revealed no outlier samples. Data were normalized to log(counts per million) with edgeR (Robinson et al., 2010) and fit to a linear model with moderated t-statistics using limma-trend (Smyth, 2005). We applied limma’s eBayes() function to identify genes that were differentially expressed across any of the four groups (high-playing males, low-playing males, high-playing females, low-playing females), as well as in specific post-hoc contrasts for the effects of play and sex. These analyses identified 4,261 genes with nominally significant differential expression in any contrast related to sex and play (*p* < 0.05). We used Weighted Gene Co-expression Network Analysis (Langfelder & Horvath, 2008), implemented with the WGCNA R package, to cluster these 4,261 genes into gene co-expression modules and to characterize their patterns of expression across the four groups. Network reconstruction was performed using the blockwiseModules() function with power = 12, corType = ‘bicor’, maxPOutliers = 0.1, networkType = ‘signed’, minModuleSize = 10, reassignThreshold = 0, mergeCutHeight = 0.25, and otherwise default parameters. This function starts with pairwise Pearson correlations among gene pairs, converts them to a signed topological overlap matrix using a power function to create an approximately scale-free edge-weight distribution, followed by hierarchical clustering with a dynamic tree-cutting algorithm. We required a minimal module size of 10, and, after initial module detection, merged any pairs of modules whose eigengenes were correlated (r > 0.75). Each module is summarized by its first principal component, termed the module eigengene. We fit the module eigengenes to a linear model with limma and calculated associations of module eigengenes with post-hoc contrasts related to sex and play. As a secondary analysis, we also considered quantitative effects of play in each sex, again using linear models implemented in limma separately by sex. We conducted enrichment analysis to identify gene ontology (GO) terms statistically overrepresented (adjusted *p*-value < 0.05) within modules of interest using the ShinyGO (V0.80; Ge, Jung & Yao, 2020).

### Single-cell RNA-sequencing (scRNA-seq)

On P4, neonates of both sexes (n=6) were deeply anesthetized with Fatal Plus and decapitated. Brains were removed and placed on ice, and the amygdala of both hemispheres was dissected out and placed in Hibernate A solution (Thermo Fisher Scientific, Waltham, MA) while the remaining brains were dissected. Single cells were isolated for transcriptional profiling by proteolytic enzymatic digestion using the Worthington Papain Dissociation System (Worthington Biochemical Corporation, Lakewood, NJ), following the manufacturer’s protocol. Samples were centrifuged, then resuspended in 0.05% bovine serum albumin in phosphate-buffered saline. Cell suspensions were then pooled, resulting in four final samples prepared for submission: 2 male samples and 2 female samples, each containing cells from 3 animals per sex. Pooled suspensions were counted using a hemocytometer and delivered to Maryland Genomics for further processing using the 10x Genomics system. A total of 3,000 cells per sample were loaded by GRC staff into each well of a chromium microfluidics container (10x Genomics, Pleasanton, CA). Sequencing libraries were prepared using the Chromium Next GEM Single Cell 3’ Kit v2 per the manufacturer’s instructions. Samples were sequenced across 4 lanes (1 sample per channel) using the 75 base pair paired-end protocol on an Illumina HiSeq4000 sequencer.

### scRNA-seq analysis

The raw scRNA-seq data were processed to counts of unique molecular identifers (UMIs) in each droplet using CellRanger, then passed through SoupX to reduce artifacts from ambient RNA (Young & Behjati, 2020). Quality control, normalization, and analysis were performed using the Seurat (v2) Standard Workflow. Cells were clustered using the Louvain method, revealing 8 major cell type clusters from which we assessed sex differences in gene expression. Iterative sub-clustering revealed 11 inhibitory neuron sub-populations. Marker genes enriched in each cell type compared to other cell types, as well as sex differences in cells from male vs. female animals within each cell type, were assessed using Wilcoxon signed-rank tests, implemented with the Seurat FindMarkers() function. We integrated our scRNA-seq and bulk RNA-seq datasets using gene co-expression networks. First, we used WGCNA to identify gene co-expression modules in the scRNA-seq data, considering the 10,517 genes expressed in more than 1% of cells. The count matrix was smoothed using KNN smoothing (k=15, d=30) and normalized using the NormalizeData() function in Seurat. This normalized matrix was then used to identify WGCNA modules, using a soft power of 4 for modularization, revealing 25 gene co-expression modules. We then assessed the distribution of these 25 modules across our identified scRNA-seq major cell types to gain insight on potential cell-type specificity of each identified module. To integrate this analysis with our bulk mRNA-seq data, we assessed the association of each scRNA-seq module with sex and play via expression of the module eigengene and compared the overlap between each scRNA-seq and bulk RNA-seq module using Fisher’s exact test.

### *In situ* hybridization

To identify play-active cells, we performed *in situ* hybridization of activity-related genes in juvenile rats following a time-monitored “play pulse” ahead of tissue collection. Juveniles (P28) of both sexes were first isolated for 2 hours to increase social motivation. Then, they were placed in an enclosure (49 x 37 cm, 24 cm high) with TEK-Fresh cellulose bedding (Envigo) with a novel same-sex, same-age, non-sibling play partner and allowed to play for 10 min. One hour later, rats were deeply anesthetized with Fatal Plus and transcardially perfused with phosphate-buffered saline (PBS; 0.1M, pH 7.4) followed by 4% paraformaldehyde (PFA; 4% in PBS, pH 7.2). Brains were removed and postfixed for 24 h in 4% PFA at room temperature, then placed in 30% sucrose solution at 4°C. When fully submerged, brains were immediately flash frozen in powdered dry ice and stored at −80°C until sectioning. Coronal sections were cut at a thickness of 20 μm on a Leica CM2050S cryostat and directly mounted onto silane-coated slides.

*In situ* hybridization was carried out using the RNAscope® multiplex fluorescence assay kit (Advanced Cell Diagnostics, Newark, CA), following the manufacturer’s instructions with small modifications. The following probes were used: Rn-Egr1 (catalog number 318571), Rn-Slc17a6 (317011-C2 and 317011-C4), Rn-Slc32a1 (424541-C3), Rn-Spen (842581-C4), Rn-Klhdc8a (842571-C4), and Rn-Cyp19a1 (520461-C2). Briefly, tissue was baked onto slides using the HybEZ^TM^ II Oven at 60°C for 30 min, then dehydrated using four 5 min dehydration steps in 50%, 75%, 100%, and 100% ethanol, respectively. Tissue was then treated with peroxidase solution for 10 min at room temperature, followed by a washing step with room temperature water. Slides were next placed in a 95°C water solution to acclimate for 10 seconds before being placed in 95°C target retrieval reagent solution for 5 min, followed by a 3 min incubation in 100% ethanol at room temperature. Slides were air-dried for 5min, then digested with Protease III solution for 30 min at 40°C in the HybEZ^TM^ II Oven. Following two 2 min washes with the supplied wash buffer, sections were incubated (again at 40°C) for 30 min in a mixed probe solution containing all necessary probes for the experiment (1:50 C2-C4 probe dilution using C1 probe solution). Slides were then washed twice and stored in 5X SSC overnight. All subsequent steps were performed at 40°C in the HybEZ^TM^ II Oven and separated by two 2 min washes. AMP1 and AMP2 solutions were added for 30 min, and AMP3 was added for 15 min. HRP C1 was added for 15 min, then the appropriate Opal dye (520, 570, 620, or 690; Akoya Biosciences, Marlborough, MA) for 30 min, followed by HRP blocker for 15 min. This procedure was then repeated consecutively for C2, C3, and C4, with a new Opal dye used for each probe channel. Following the RNAscope® procedure, slides were stained immediately with Hoescht (1:2000; Invitrogen, Waltham, MA) for 5 min and cover-slipped using Prolong Diamond Anti-Fade Mountant (Thermo Fisher Scientific, Waltham, MA). Slides were imaged on a Nikon CSU-W1 microscope equipped with 405, 488, 561, and 647 lasers, using a 20x (0.75 NA) and 60x (1.49 NA) TIRF oil-immersion objective, and analyzed using Imaris (Bitplane, Zurich, Switzerland) software.

## Results

### The transcriptome of playfulness in the MeA of juvenile rats is sex-biased

To assess play-associated gene expression patterns and whether these differ between the sexes, we first quantified play behaviors in a cohort of n=18 male and n=18 female juvenile rats derived from 4 litters. Animals were weaned on P21 and allowed to play with a sex- and age-matched non-sibling play partner once daily for 10 minutes on P26-29 (Figure 1a). As expected, we observed a sex difference in play, with males exhibiting a higher total (i.e., cumulative) play score on average than females (*p* = 0.04; Figure 1b). Animals with play scores within the top third per sex were designated as “high-playing” and those with scores in the bottom third per sex were designated as “low-playing”, resulting in four groups of 6 animals each: male high-playing (MHI), male low-playing (MLO), female high-playing (FHI), and female low-playing (FLO). Animals with mid-range play scores were excluded from further analyses with the expectation that the greatest differences in gene expression patterns would be observed in animals on the extremes. Subjects were sacrificed on P30, and MeA tissue was collected and processed for bulk RNA-sequencing.

**Figure 1.**
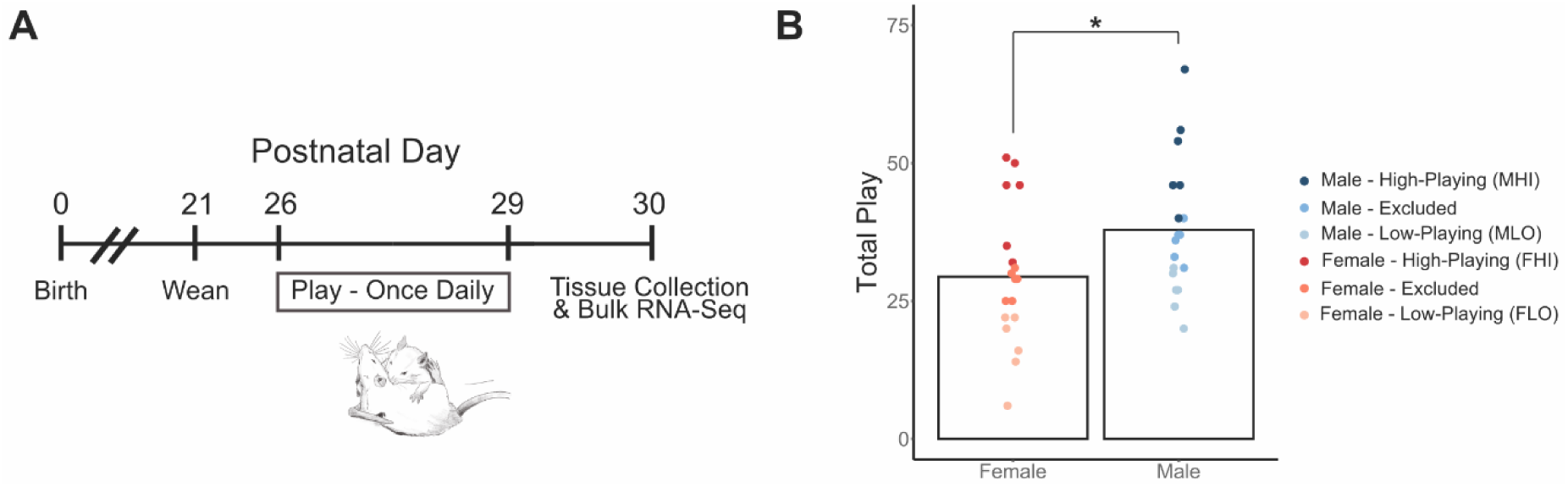
Framework for bulk RNA-sequencing experiment. (***a***) Timeline of experimental procedures. (***b***) Quantification of cumulative number of play behaviors following four days of testing, used to determine high- and low-players for later RNA-sequencing, with the top third per sex designated as “high-playing” and bottom third per sex as “low-playing”. Bars indicate group means ± SEM, and points represent data from individual rats. \**p* < 0.05, *n* = 6 per group.

Analysis of the bulk RNA-seq data revealed 4,261 genes with nominal differential expression (*p* < 0.05) related to sex or play. We then used Weighted Gene Co-Expression Network Analysis (WGCNA; Zhang & Horvath, 2005; van Dam et al., 2018), to cluster these 4,261 genes into 22 gene co-expression modules based on their coordinated transcriptional patterns (Figure 2a, Figure S1, Table S1). Many of these gene networks showed a strong association with social play based on expression patterns of the module eigengene, or the first principal component of the genes in the network. Statistical associations of modules with play were computed using linear models, with effects of play in each sex determined using post-hoc contrasts. We also considered quantitative effects of play in each sex, again using linear models implemented separately by sex.

**Figure 2.**
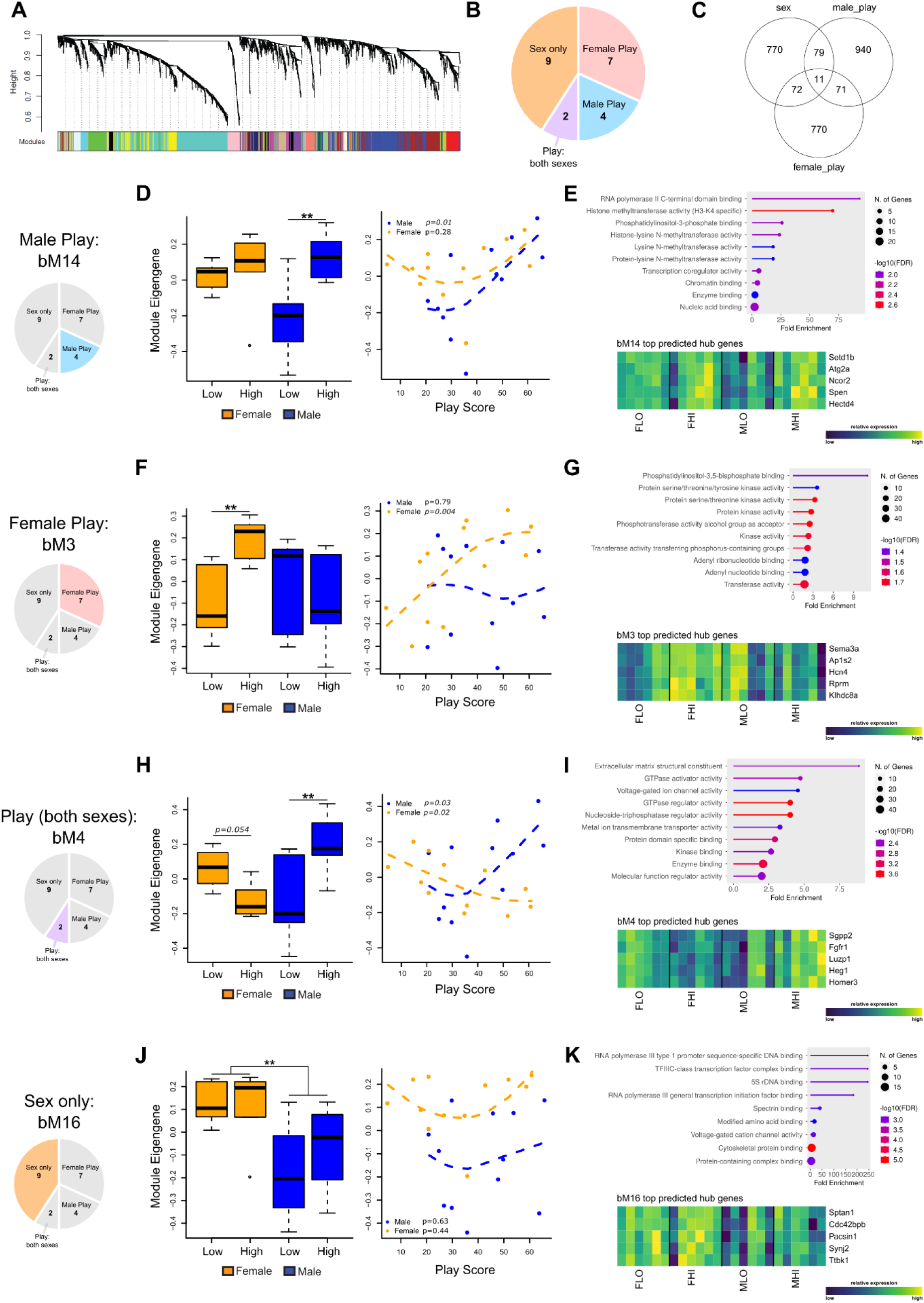
The transcriptome of playfulness in the juvenile rat medial amygdala is sex-biased. (***a***) Cluster dendrogram of the 4,261 genes used for the weighted gene co-expression network analysis (WGCNA), with each color representing one of the 22 modules identified. (***b***) Pie chart showing the number of bulk WGCNA modules in each category: modules associated with playfulness in females (“Female Play”), playfulness in males (“Male Play”), playfulness in both sexes (“Play: both sexes”), or those that only showed an association with sex, not playfulness (“Sex only”). See Figure S1 and Table S2 for graphs and statistics for all 22 modules. (***c***) Venn diagram showing the number of differentially expressed genes (DEGs; for nominal *p* < 0.05) associated with sex (male vs. female), male playfulness (“male_play”; high-vs. low-playing males), and female playfulness (“female_play”; high-vs. low-playing females). (***d, f, h, j***) The majority of play-associated DEGs are sex-specific. Module eigengene expression across groups (left) and in relation to cumulative play score (right; y-axis is the same for both groups) for the four representative modules bM14 (Male Play), bM3 (Female Play), bM4 (Play: both sexes), and bM16 (Sex only), respectively. (***e, g, i, k***) Information on each representative module, including Gene Ontology enrichments (top) and heat maps showing expression of the top five predicted hub genes for each module across each of the 6 subjects per group (bottom). “Low” = low-playing; “High” = high-playing; FLO = female low-playing; FHI = female high-playing; MLO = male low-playing; MHO = male-high-playing. \**p* < 0.05, ** *p* < 0.01, *n* = 6 per group.

We categorized each module into one of four categories based on their statistical associations with playfulness and/or sex. Notably, only two of the 22 modules showed a statistical association (for FDR-adjusted *p* < 0.05) with playfulness in both sexes (“Play: both sexes”; Figure 2b and Table S2). Many more modules were associated with playfulness in one sex but not the other; seven modules were associated with playfulness in females but not in males (“Female Play”), while four were associated with playfulness in males but not females (“Male Play”). We designated the remaining nine modules as “Sex only”: these modules showed no significant association with playfulness but were significantly associated with sex, likely representing generalized transcriptional sex differences in the juvenile MeA. This finding – that the majority of playfulness-associated modules are associated with play level in rats of one sex but not the other – suggests that there is a distinct transcriptomic profile associated with playfulness in the male MeA compared to that of females.

This minimal overlap was also seen when assessing patterns of playfulness-associated differentially expressed genes (DEGs). We identified DEGs associated with all pairwise comparisons between groups, focusing on the sets of DEGs associated with “male play” (i.e., playfulness in males; MHI vs. MLO) or “female play” (i.e., playfulness in females; FHI vs. FLO), as well as those associated with sex in general (male vs. female). Few genes exceeded stringent false discovery rates after accounting for multiple testing. However, at a nominal (uncorrected) *p*-value < 0.05, we found 1,101 “male play” DEGs and 924 “female play” DEGs. Similar to the minimal overlap seen in playfulness-associated WGCNA modules, only 82 of these playfulness-associated DEGs were shared between the sexes (<9%; Figure 2c). The playfulness-associated gene signatures were also largely distinct from those associated with generalized sex differences: the majority of sex-biased DEGs were not shared with playfulness-associated DEGs (162 of 932 total sex-biased DEGs; 17.4%).

The identified WGCNA modules are diverse, ranging in size from 21 to 688 genes with a variety of predicted functions. Modules associated with male playfulness included bM14 (named as such as it was the 14^th^ largest module in terms of the number of genes it contains), a module which consisted of 66 genes, many of which have roles in transcriptional regulation. Gene Ontology (GO) analysis of the genes within this module highlighted terms pertaining to chromatin binding, transcription coregulator activity, and histone methyltransferase activity (Figure 2e, top). Expression of the bM14 module eigengene was significantly different between low- and high-playing males (*p* = 0.001) but not between low- and high-playing females (*p* = 0.72) and was associated with quantitative play score in males (*p* = 0.01) but not females (*p* = 0.28; Figure 2d). Identified hub genes – highly connected genes within a module predicted to serve as key driver genes, derived by the Pearson correlation between each gene and the module eigengene (“module membership” score) – for bM14 included transcriptional regulators like *Spen* and *Ncor2,* which were significantly upregulated in high-playing males compared to low-playing males (Figure 2e, bottom), following the pattern seen in the bM14 eigengene.

Of the seven modules associated with playfulness in females, bM3 showed the strongest association: eigengene expression differed significantly between low- and high-playing females (*p* = 0.0039) and was strongly associated with play score in females but not males (*p* = 0.004 and 0.79, respectively; Figure 2f). This module consists of 306 genes and is significantly enriched for GO terms suggesting a role in signal transduction, including kinase activity and transferase activity (Figure 2g, top). bM3 hub genes include *Klhdc8a*, a kelch-domain containing protein believed to be part of the MAPK signal pathway (Zhu et al., 2020) and *Hcn4*, encoding a hyperpolarization-activated, cyclic nucleotide-gated potassium channel (Figure 2g, bottom).

Two modules – bM4 and bM7 – were associated with playfulness in both sexes (Figure S1c). Strikingly, however, this was in opposite directions in males and females. This inverse relationship was seen most strongly in bM4 (Figure 2h). While expression of the module eigengene was higher in high-compared to low-playing males (*p* = 0.002) and was positively correlated with play score in males (*p* = 0.03), eigengene expression was negatively correlated with play score in females (*p* = 0.02) and lower in high-compared to low-playing females, although this effect was only trending (*p* = 0.054). Genes in this shared play module were largely associated with the cell membrane/membrane signaling, as there were significant associations with GO terms suggesting a role in voltage-gated ion channel activity, metal ion transmembrane transporter activity, and the extracellular matrix (Figure 2i, top), including hub genes like *Homer3*, a postsynaptic density scaffolding protein, and *Heg1*, a negative regular of membrane permeability (Figure 2i, bottom).

The remaining nine modules, deemed “Sex only” as they showed no association with playfulness, consisted of a few modules driven primarily by genes on the X chromosome (e.g. bM22, including its predicted hub genes *Kdm6a*, *Pbdc1,* and *Eif2s3*) or on the Y chromosome (e.g. bM21, including its predicted hub genes *Uty*, *Eif2s3y*, and *Ddx3*). Other modules, however, were made up of mostly autosomal genes and likely represent generalized transcriptional sex differences in the juvenile MeA. This included bM16, whose module eigengene showed higher expression in females (both low- and high-playing) than males (Figure 2j) and consisted of genes with roles in cytoskeletal protein and complex binding like predicted hub genes *Ttbk1*, encoding tau tubulin kinase 1, and *Sptan1*, encoding spectrin alpha (Figure 2k). While our focus in this manuscript is on playfulness-associated modules, these generalized “sex only” modules may be of interest for future studies on juvenile sex differences in gene expression in the MeA, a brain region with known roles in various socio-sexual behaviors.

### Sex-biased DEGs in the neonatal rat amygdala are enriched in inhibitory neurons

We previously described a critical period for sexual differentiation of the amygdala during the first four days of life in the rat (VanRyzin et al., 2019). In males, high androgen levels produced by the fetal testis result in a higher endocannabinoid tone in the neonatal amygdala, inducing microglia to become more phagocytic. This results in a sex difference in astrocyte density in the juvenile MeA and masculinized play (VanRyzin et al., 2019). To further explore the origins of these cell type-specific sex differences and their potential contributions to sex differences in juvenile play, we generated single-cell RNA-seq (scRNA-seq) data from the amygdala of 4-day-old male and female rats (*n* = 6 animals per sex; Figure 3a). Following quality control, we studied the transcriptomes of 11,756 cells. Louvain clustering identified nine major cell types. As expected, inhibitory neurons (IN; *Gad1*+ or *Gad2+*) and astrocytes (*Slc1a3*+) were the most abundant, followed by microglia (*Aif1*+), excitatory neurons (EN; *Slc17a6*+ or *Slc17a7+*), oligodendrocyte precursor cells (OPC; *Pdgfra*+), endothelial cells (*Flt1+*), fibroblasts (*Dcn+*), ependymal cells (Mif1+), and mural cells (*Rgs5*+) (Figure 3a-3b; Figure S2; Table S3).

**Figure 3.**
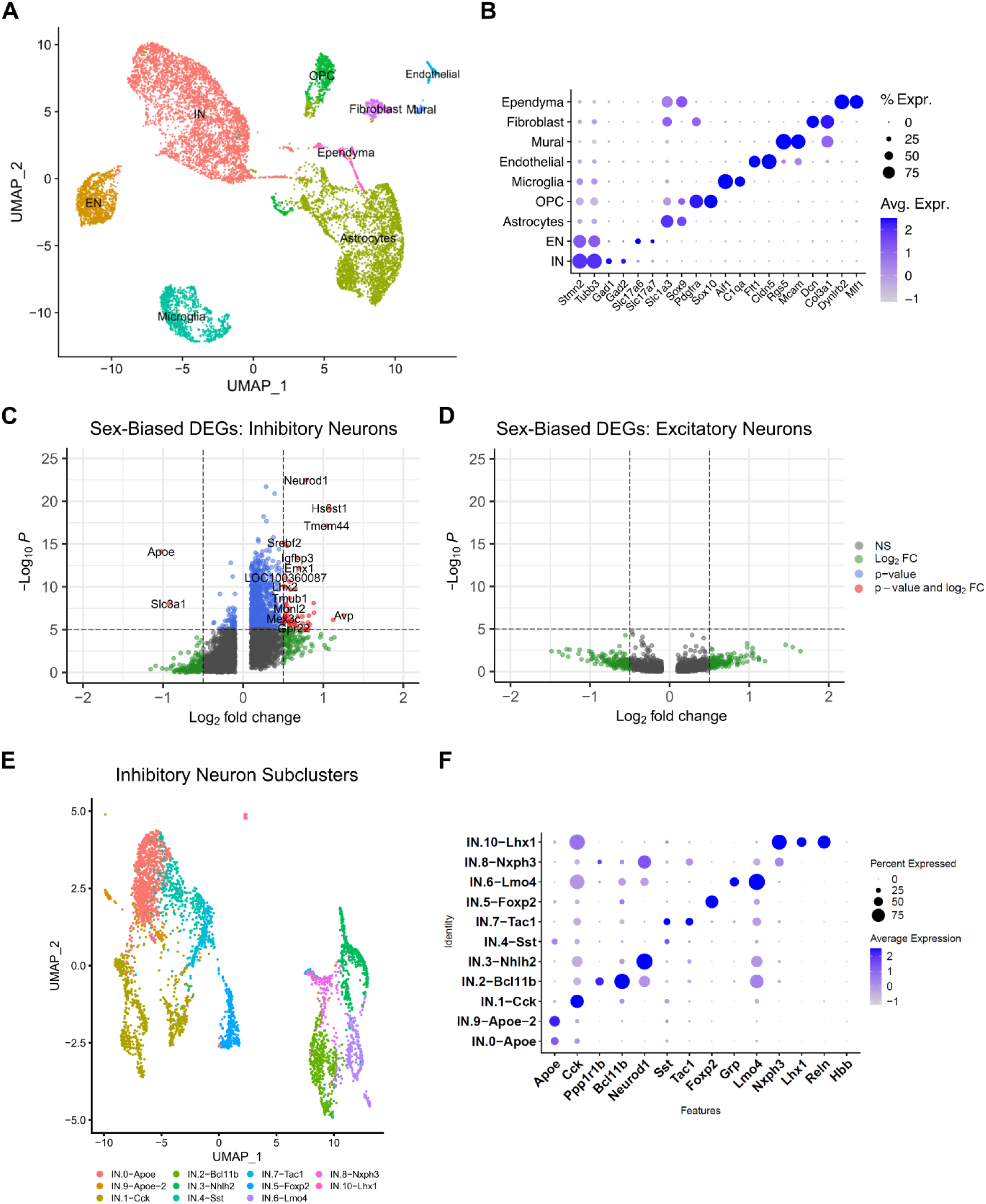
Single-cell RNA-sequencing analysis indicates that there are more sex-biased DEGs in inhibitory neurons than excitatory. (*a*) UMAP plot identifying 9 major cell types in the neonatal amygdala of 2 male and 2 female samples, each containing pooled samples from 3 pups each. IN = inhibitory neurons, EN = excitatory neurons, OPC = oligodendrocyte precursor cells. (*b*) Dot plot showing the expression of marker genes used to identify the 9 major cell types. (*c* and *d*) Volcano plots showing sex-biased DEGs (male-biased: log_2_ fold change > 0) within inhibitory neurons (*c*) and excitatory neurons (*d*) with shared legend to the right of *d*. Genes colored in red showed significant sex-biased expression with both a significant *p*-value as well as log_2_ fold change. (*e*) Subclustering of the inhibitory neuron population, resulting in 11 distinct subtypes (IN.0 – IN.11), with the marker genes differentiating these subtypes shown in *f*.

There were no sex differences in the proportions of major cell types in males compared to females. However, clear sex differences emerged when we assessed sex-biased DEGs (genes showing differential expression between males and females) within the identified major cell types. Notably, the overwhelming majority of sex-biased DEGs were found in the inhibitory neuron population (727 DEGs at FDR < 0.01; Figure 3c; Figure S3; Table S4), while relatively few were found in the excitatory neuron population (35 DEGs in all other cell types combined; Figure 3d). Down-sampling analyses confirmed that this pattern was not explained by inhibitory neurons being more abundant than other cell types. Calculating the effect sizes by two analytical approaches demonstrated consistent effects (Figure S4; Table S5). This finding – that GABAergic neurons exhibit greater transcriptional sex differences than glutamatergic neurons in the developing rat amygdala – is consistent with that seen in a scRNA-seq analysis of sex-biased DEGs in MeA of adult mice (Chen et al., 2020), suggesting that the sex-biased gene expression patterns that we observe early in life are long-lasting.

Sex-biased inhibitory neuron DEGs included *Neurod1* (log2(foldchange) in males vs. females = 0.79; p-value = 3.5e-23; Figure S5), which encodes a transcription factor that functions in neuronal differentiation and plasticity (Tutukova et al., 2021) and *Crym* (logFC = 0.33, p-value = 6.1e-11), which encodes a thyroid hormone-binding protein previously identified as a critical modulator of the sex-specific response to social isolation in the adolescent MeA (Walker et al., 2022). While there were fewer DEGs within excitatory neurons, these included *Cnr1* (logFC = 1.0, p-value = 1.8e-3), encoding cannabinoid receptor 1, which was more highly expressed in males, an interesting finding given our previous discovery that a higher endocannabinoid tone in males induces higher microglial phagocytosis in the developing amygdala (VanRyzin et al., 2019). Although many neurons are not fully differentiated at this neonatal timepoint, subclustering revealed 11 distinct classes of inhibitory neurons (Figure 3e; Table S6), many of which specifically expressed canonical markers such as neuropeptides and transcription factors (e.g. *Sst*, *Cck*, and *Foxp2*; Figure 3f).

### Integration of bulk and scRNA-seq analyses confirms and expands “male play” and “female play” modules

To better understand potential cell-type specificity, key driver genes, and predicted functions of our 22 identified play-associated gene modules, we developed a network analysis approach to integrate our scRNA-seq and bulk RNA-seq analyses. Using similar procedures as for our bulk RNA-seq dataset, we conducted WGCNA on the scRNA-seq dataset, identifying 25 gene co-expression modules (Figure 4a; Table S7). Several of these modules were expressed primarily in a single cell type. For instance, we identified modules enriched in microglia (scM3: 894 genes), inhibitory neurons (scM4, scM5, and scM6: 2,376 genes combined), and ependymal cells (scM7: 683 genes), as shown in Figure 4b.

**Figure 4.**
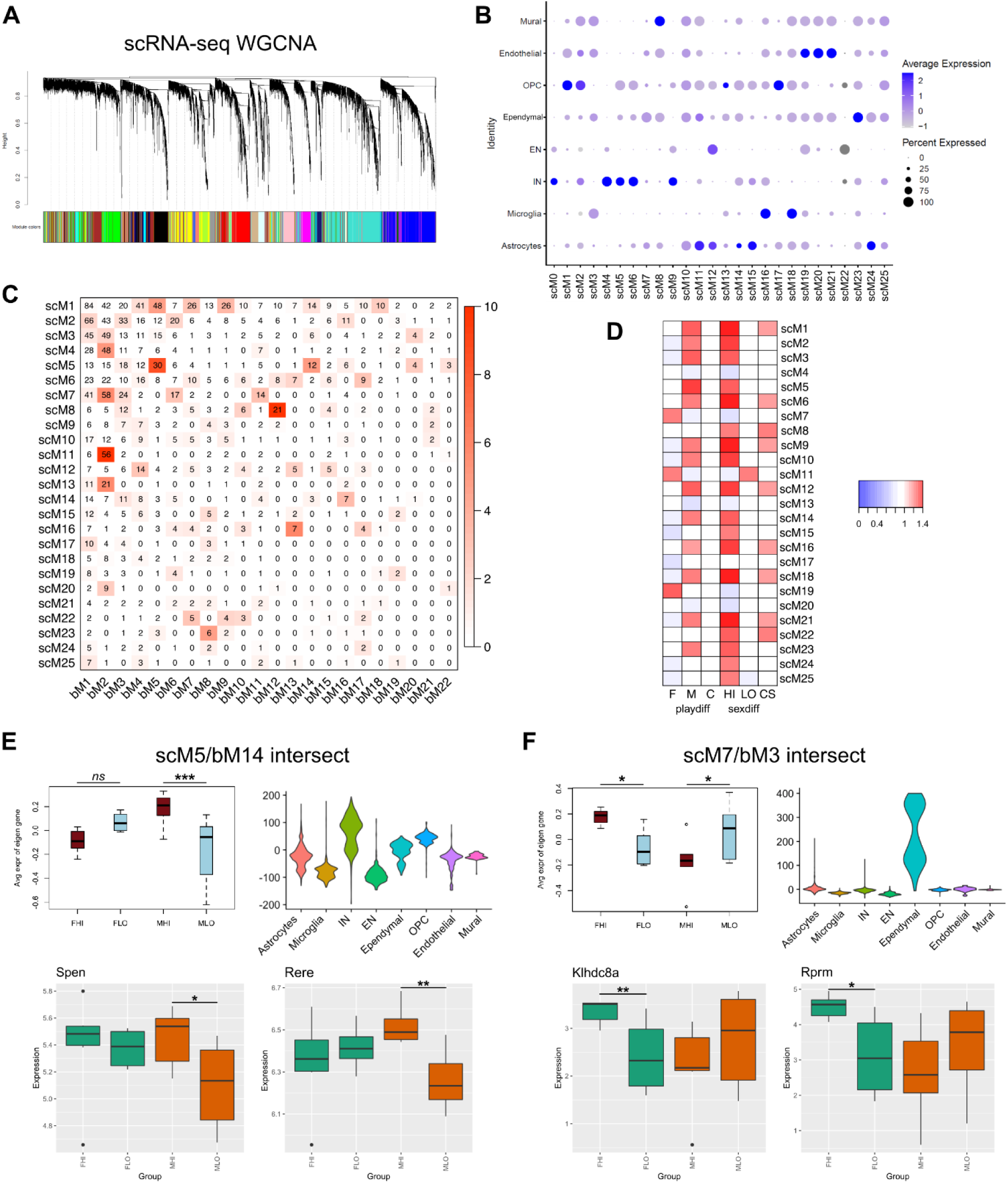
Integration of WGCNA analyses identifies predicted cell-type specificity and likely hub genes for selected modules. (***a***) Cluster dendrogram from the network reconstruction of the scRNA-seq data, using the set of 4,261 genes from the bulk-seq WGCNA as input, with each color representing one of the 25 modules identified. (***b***) Dot plot showing likely major cell type identity of each of the 25 scRNA-seq modules. (***c***) Matrix showing the number of overlapping genes between each single-cell (scM1-scM25) and bulk-seq (bM1-bM22) module, with color indicating the −log_10_(*p*-value) of a Fisher’s exact test. (***d***) Heat map showing module-trait relationships (using module eigengene) across various comparisons: F = (FHI-FLO), M = (MHI-MLO), C = (MHI+FHI)/2-(MLO+FLO)/2, HI = (MHI-FHI), LO = (MLO-FLO), CS = (MHI+MLO)/2-(FHI+FLO)/2. (***e*** and ***f***) Further interrogation of the two representative intersecting module sets, including plots showing the average single-cell module eigengene across the 4 play groups (top left), association with major cell type categories (top right), and box plots showing expression of the 2 top-predicted candidate hub genes (bottom): those genes within the set of shared genes common to scM5 & bM14 (***e***) and scM7 & bM3 (***f***) that had the top-2 highest intramodular connectivity within the bulk-seq dataset. IN = inhibitory neurons, EN = excitatory neurons, OPC = oligodendrocyte precursor cells, FHI = female high-playing, FLO = female low-playing, MHI = male high-playing, MLO = male low-playing. \**p* < 0.05, ** *p* < 0.01, *** *p* < 0.001, *n* = 6 per group.

Providing confidence in our analyses, multiple scRNA-seq modules showed significant overlap with the genes in particular bulk RNA-seq modules (Figure 4c; Table S8). This included scM11, a module enriched in astrocytes, which showed significant (*p* < 0.001) overlap with bM2, a bulk RNA-seq module associated with male play, and scM16, a module enriched in microglia, which significantly (*p* < 0.001) overlapped with the female play module bM13. We next projected the eigengenes of scRNA-seq-derived modules onto our bulk RNA-seq samples to assess their associations with sex and play. As expected given the sex-biased expression of the playfulness-associated bulk RNA-seq modules previously described (Figure 2), most scRNA-seq modules were associated with playfulness in only one sex (Figure 4d and Table S9). Many scRNA-seq modules showed significant associations with playfulness in males but not in females based on expression of the module eigengene, with scM1, scM5, and scM12 exhibiting especially strong associations, while scM7, scM11, and scM19 showed strong associations with playfulness in females but not males.

We next selected two representative bulk RNA-seq modules to explore in greater detail. This included the representative “male play” module bM14, a module that, as previously described (Figure 2d-e), is associated with playfulness in males but not females and consists of many genes involved in transcriptional regulation. bM14 shows significant overlap with the genes in scRNA-seq module 5 (scM5; *p* < 0.001) with 12 genes shared between the two (Figure 4c; Table S8). scM5, via the module eigengene, also shows significant association with playfulness in males but not females (*p* < 0.001; Figure 4e, top left panel) and is enriched in inhibitory neurons (Figure 4e, top right panel), a notable result given sex-biased DEGs are enriched within inhibitory compared to excitatory neurons within this same dataset (Figure 3c). Within the set of 12 genes shared between bM14 and scM5, we noted multiple genes that were also ranked highly as hub genes for bM14 – those with a high “module membership” (MM) score indicating a strong degree of intramodular connectivity, suggesting a key regulatory role. This included *Spen*, (spen family transcriptional repressor; bM14 MM = 0.909) and *Rere* (arginine-glutamic acid dipeptide repeats; bM14 MM = 0.854). Notably, both of these genes have known roles in nuclear receptor-mediated transcription, and deleterious variants in both genes are associated with neurodevelopmental disorders, including autism spectrum disorder (Radio et al., 2021; Niehaus et al., 2022). Expression of each of these predicted hub genes is also associated with playfulness in males but not females (*p* = 0.032 and 0.008 for *Spen* and *Rere*, respectively; Figure 4e, bottom).

We also further explored representative “female play” module bM3, a bulk RNA-seq module associated with playfulness in females but not males that is enriched for GO terms suggesting a role in signal transduction (Figure 2f-g). bM3 shows significant overlap with scRNA-seq module 7 (scM7; *p* = 0.014), with 24 genes shared between the two (Figure 4d; Table S8). Interestingly, while bM3 is specifically associated with playfulness in females, but not males, scM7 is associated with playfulness in both sexes, but in opposite directions (coincidentally, *p* = 0.014 for both male play and female play). Expression of the scM7 eigengene is higher in high-playing females than in low-playing females, while this pattern is reversed in high- and low-playing males (Figure 4f, top left panel). Analysis as to the potential cell-type specificity of scM7 suggests that genes in this module are enriched in ependymal cells (Figure 4f, top right panel), a type of epithelial glial cell. From the list of 24 genes shared between bM3 and scM7, genes with the highest module membership score (indicating a high-confidence prediction as potential hub genes) within bM3 included *Klhdc8a* (kelch domain containing 8a) and *Rprm* (reprimo), both of which showed an association with playfulness in females but not males (*p* = 0.008 and 0.013 for *Klhdc8a* and *Rprm*, respectively; Figure 4f, bottom). While little is known about *Klhdc8a*’s role in the brain, a recent mouse study showed that expression of *Rprm* is female-biased in the adult hypothalamus, where it plays a role in estrogen receptor-dependent regulation of energy expenditure (van Veen et al., 2020).

### The majority of play-active cells in the juvenile MePD are inhibitory neurons, with more in males than females

By this point in our analyses, two pieces of evidence – that the majority of sex-biased DEGs are within inhibitory neurons in the neonatal amygdala (Figure 3c), and that expression of one of our major play-associated modules of interest (bM14) appears enriched within this same cell type (Figure 4e) – suggested that inhibitory neurons in the MeA may represent an important cell type driving sex differences in social play. To test this hypothesis, we designed an *in situ* hybridization experiment using fluorescent RNAscope®. Juvenile male and female rats were sacrificed one hour after a playful experience with a novel play partner. Tissue sections containing MeA were then processed for RNAscope®, using probes for *Vgat* (inhibitory neuron marker), *Vglut2* (excitatory neuron marker), and the immediate early gene *Egr1* (also known as *Zif268*), to label play-active cells (Figure 5a-b). We also included a probe for *Cyp19a1* (aromatase), following recent work that demonstrated that aromatase-expressing cells in the MeA are critical for various social behaviors that are sexually differentiated (Unger et al., 2015; Yao et al., 2017; Figure 5b).

**Figure 5.**
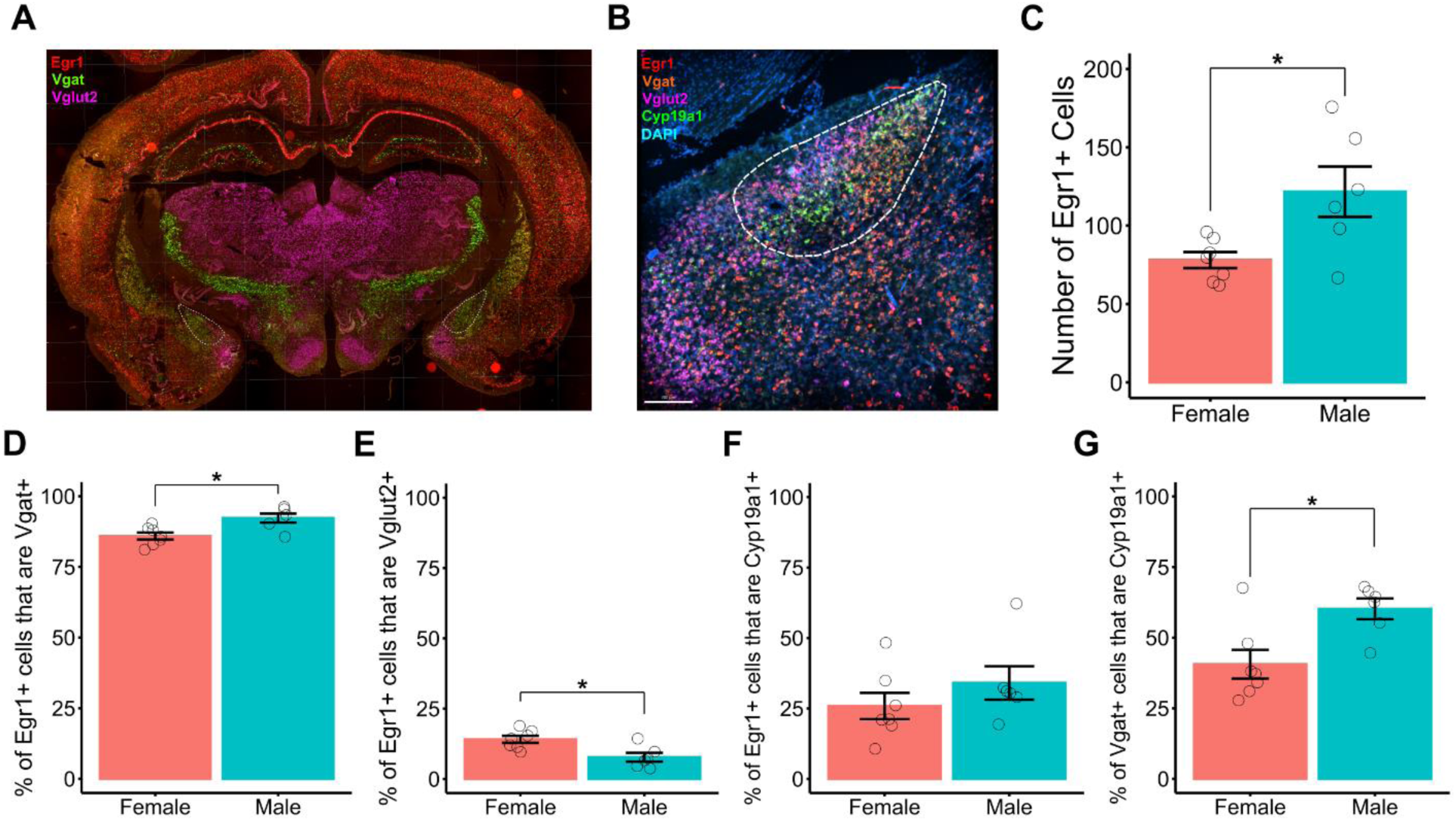
RNAscope® phenotyping of play-active cells. (***a***) Representative low-magnification image of *Egr1* (red), *Vgat* (green), and *Vglut2* (fuschia) expression via fluorescent *in situ* hybridization (RNAscope). White dashed outline indicates the posterodorsal medial amygdala (MePD) (***b***) Representative high-magnification image of *Egr1* (red), *Vgat* (orange), *Vglut2* (fuschia), and *Cyp19a1* (green) expression alongside DAPI (blue). White dashed outline indicates the MePD. (***c***) Quantification of the number of play-active (Egr1+) cells in the male and female MePD. The percentage of Egr1+ cells co-expressing *Vgat* (***d***) and *Vglut2* (***e***) was also calculated, as well as the percentage of Egr1+ cells (***f***) and Vgat+ cells (***g***) that co-expressed *Cyp19a1* (aromatase). Bars indicate group means ± SEM, and open circles represent data from individual rats. \**p* < 0.05; *n* = 6-7 per group.

We focused our analyses on the posterodorsal MeA (MePD), as we had previously demonstrated that subregion is the critical node determining MeA-driven sex differences in play (VanRyzin et al., 2019). Replicating our previous work using immunohistochemistry (VanRyzin et al., 2019), males exhibited significantly more play-active (Egr1+) cells in the MePD (*p* = 0.04; Figure 5c). In both sexes, the vast majority of play-active cells in the MePD were GABAergic (∼90% Vgat+ vs. ∼10% Vglut2+); however, the distribution between inhibitory and excitatory neurons significantly differed between males and females (Figure 5d-e). In the male MePD, 92% of play-active cells were GABAergic, with the remaining ∼7% co-labeling with *Vglut2*. In females, the percentage of glutamatergic play-active cells was twice that seen in males: 14% of Egr1+ cells were glutamatergic, while 86% were GABAergic, a significant difference from that seen in males (*p* = 0.01). Additionally, around a third of play-active cells expressed aromatase (Figure 5f). While there was no sex difference in the percentage of Egr1+ cells that were Cyp19a1+ (*p* = 0.30), there was a significant sex difference in the proportion of GABAergic cells that expressed aromatase in this region, with 60% of GABAergic cells in males co-labeling with *Cyp19a1*, while only 40% of GABAergic cells did so in females (*p* = 0.01; Figure 5g).

### RNAscope® phenotyping of cells expressing key hub genes *Spen* (bM14) and *Klhdc8a* (bM3)

As our previous findings highlighted the involvement of MeA inhibitory neurons in sex differences in play, we wondered if expression of key hub genes in our selected “male play” (bM14) and “female play” (bM3) modules was also enriched within this population. We assessed expression of *Spen* and *Klhdc8a* – top-predicted hub genes for bM14 and bM3, respectively – across the major cell types identified in our scRNA-seq dataset (Figure 3b). *Spen* was expressed across multiple cell types in the neonatal amygdala, including a subset of inhibitory neurons (Figure 6a). *Klhdc8a* was expressed specifically in a subset of inhibitory neurons (log2FC = 1.9; p-value = 4.0e-107; Figure 6b).

**Figure 6.**
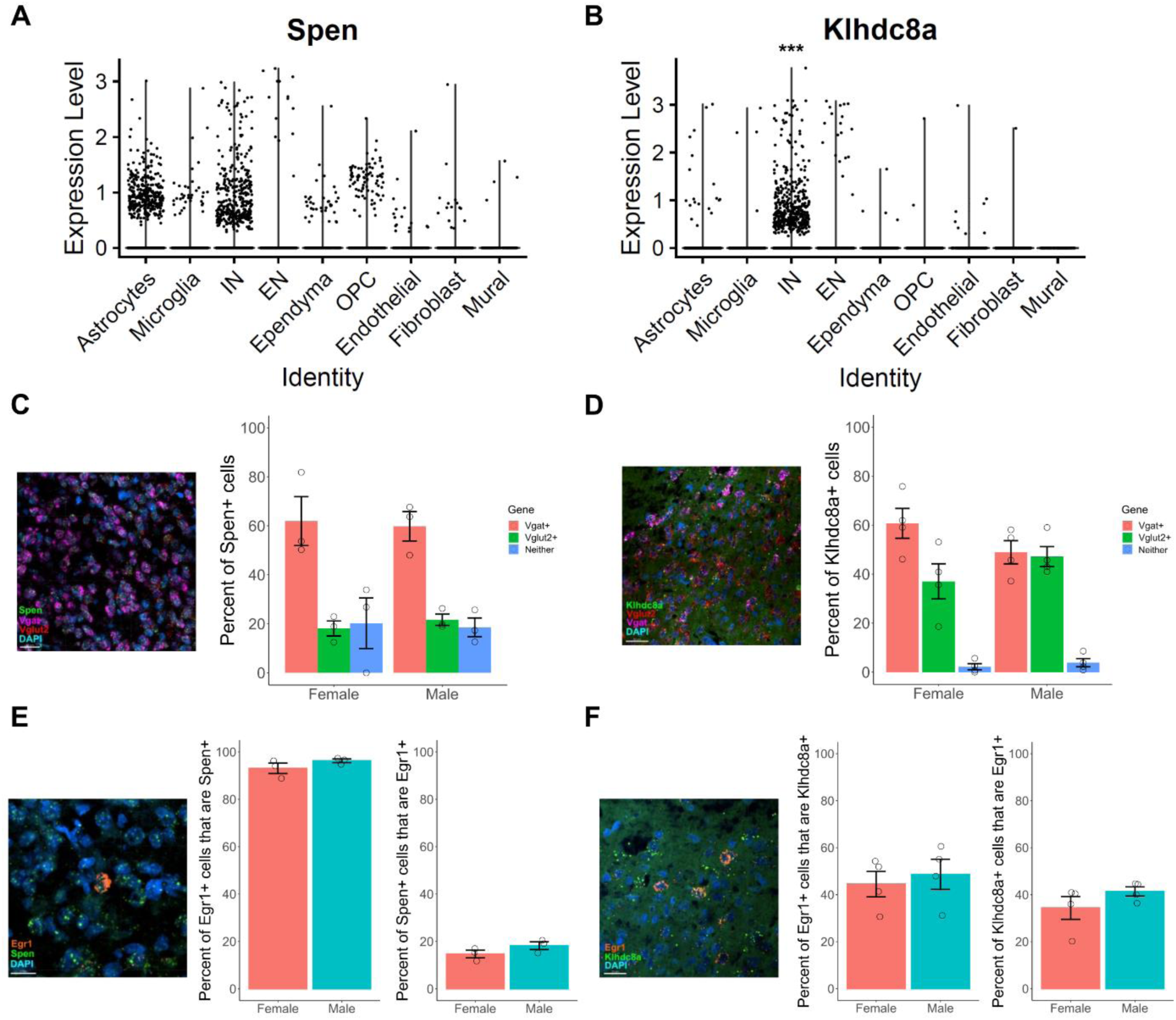
Key module hub genes *Spen* and *Klhdc8a* are enriched in the inhibitory neuron population. (***a*** and ***b***) Violin plots showing log-normalized expression of *Spen* and *Klhdc8a*, respectively, across major cell types in the neonatal rat amygdala as identified in the scRNA-seq dataset. (***c*** and ***d***) Left, representative images of *Spen* (***c***) and *Klhdc8a* (***d***) (green), *Vglut2* (red), and *Vgat* (fuschia) expression via fluorescent *in situ* hybridization (RNAscope®), alongside DAPI (blue). Right, quantification of the percentage of Spen**+** (***c***) and Klhdc8a*+* (***d***) cells that co-expressed *Vgat*, *Vglut2*, or neither neuronal marker in the male and female MePD. (***e*** and ***f***) Left, representative images of *Spen* (***e***) and *Klhdc8a* (***f***) (green) and *Egr1* (orange) expression alongside DAPI (blue). Middle, quantification of the percentage of Egr1+ cells that co-expressed *Spen* (***e***) and *Klhdc8a* (***f***). Right, quantification of the percentage of Spen+ (***e***) and Klhdc8a+ (***f***) cells that co-expressed *Egr1*. Bars indicate group means ± SEM, and open circles represent data from individual rats. *n* = 3-4 per group. IN = inhibitory neurons, EN = excitatory neurons, OPC = oligodendrocyte precursor cells.

To validate these findings and assess whether there are differences in the number of inhibitory vs. excitatory neurons expressing these hub genes in the juvenile MeA, we turned back to RNAscope®. Using alternate sections from the same set of tissue used in the prior experiment, we conducted RNAscope® using probes for *Vgat, Vglut2*, and either *Spen* or *Klhdc8a*. We also assessed whether *Spen* and *Klhdc8a* were enriched among cells activated during a play bout, using *Egr1* to mark play-active cells as before. Following our predictions, the majority of Spen+ cells (around 60%) were GABAergic in both sexes (Figure 6c). Around 20% were glutamatergic, while the remaining 20% did not co-label with either marker, suggesting they represent non-neuronal Spen+ cells. Notably, nearly all (∼95%) play-active (Egr1+) cells expressed *Spen* (Figure 6e, middle panel). There were no sex differences observed in any of these measures for *Spen*.

Supporting the results seen in the scRNA-seq enrichment analysis (Figure 6b), the majority of Klhdc8a+ cells were GABAergic (Figure 6d). In addition, there was a trending sex difference in the relative proportions of GABAergic vs. glutamatergic Klhdc8a+ cells in males compared to females (*p* = 0.087 for chi-square test of independence). In males, Klhdc8a+ cells were evenly distributed amongst excitatory and inhibitory neuron populations: ∼49% of Klhdc8a+ cells co-labeled with *Vgat,* while 47% were Vglut2+ (Figure 6d). In females, however, the balance was shifted more towards the inhibitory neuron population: 61% of Klhdc8a+ cells co-labeled with *Vgat*, while 37% co-labeled with *Vglut2*. While not the majority, a sizable percentage of Klhdc8a+ cells were play-active: ∼37% co-labeled with *Egr1* (Figure 6f, right panel). Conversely, ∼46% of play-active cells were Klhdc8a+, with no sex differences observed on either measure. Together, these studies further clarify the phenotype of cells expressing our hub genes of interest and provide additional support to the notion that MeA inhibitory cells play a central role in the establishment of sex differences in social play.

## Discussion

Play is a transient, complex social behavior that uniquely intersects with motivation and reward, motor learning, social memory and sex differences. Although play is broadly exhibited across mammalian species and has strong face validity in humans, it is remarkably poorly understood. This is likely a byproduct of the fact that inbred laboratory mice do not exhibit complex play (Pellis & Pellis, 2017a; Dvorzhak et al., 2024), thus limiting the ability of scholars of play to apply advanced genomic and molecular tools. To our knowledge, only three studies have previously assessed gene expression patterns associated with social play using RNA-seq: one profiling the amygdala (Alugubelly et al., 2019) and two profiling the medial preoptic area (Zhao et al., 2020; Polzin et al., 2024). However, these studies only included male subjects and therefore could not assess the impact of sex on play-associated gene expression – a variable which, as we demonstrate here, has a substantial influence on the transcriptional patterns observed.

We determined that play-associated gene networks in the juvenile rat are largely distinct between the sexes, with unique transcriptional modules associated with play in males and in females. This finding – that the transcriptomic landscape associated with play is qualitatively different in males and females – is especially noteworthy given the common hypothesis that play functions to shape neural circuitry enabling expression of adult behavior, an idea supported by play deprivation studies. As the MeA is fundamental to the expression of adult behaviors with known sex-differential patterns, such as parenting (Chen et al., 2019) and aggression (Edwards & Rowe, 1975; Haller, 2018), the play-associated gene networks we identified may be sex-specific for a reason: because play is serving to modulate MeA circuitry in a sex-biased way to enable such sex-typical behavior later in life. Indeed, in our previous behavioral studies (Marquardt et al., 2023), juvenile play deprivation resulted in sex-biased deficits in various socio-sexual behaviors in adulthood. Adult males deprived of play as juveniles exhibited decreased sexual behavior, hypersociability, and increased aggressiveness, with no effects seen on these measures in females, although maternal aggression was not assessed. Our identified sex-biased, playfulness-associated modules may thus represent patterns of genes, activated by the experience of play, that sculpt sex-biased maturation of the MeA. Blocking this experience-dependent, sex-biased gene activation during the “critical period” of the juvenile play window may underlie the behavioral deficits we observed in adults deprived of play as juveniles.

Indeed, previous studies have demonstrated that the experience of social play (or the lack thereof) can induce plastic changes in brain structure and function. For example, neurons in the medial prefrontal cortex of play-deprived rats exhibit longer and more branched dendrites than control animals allowed to play in their home cage (Bell et al., 2010). Conversely, neurons in the orbitofrontal cortex are more complex in rats allowed to play with multiple partners as compared to play-deprived rats or those only given access to a single partner (Bell et al., 2010). A similar phenotype was found in the anterodorsal MeA: dendrites of neurons in this region were significantly more branched in play-deprived rats compared to controls (Cooke & Shukla, 2011), suggesting that participating in play is critical for appropriate dendritic arborization and patterning, or that the experience of social isolation may induce these phenotypes. Similarly, play experience induces growth factor expression in the amygdala and various cortical regions (Gordon et al., 2003; Burgdorf et al., 2010). While these studies were only conducted in one sex (typically males) and did not assess potential sex differences, they indicate that play can indeed induce plastic changes to the juvenile brain in a potentially lasting manner. As MeA architecture and circuitry differs in adult males and females (Hines et al., 1992; Cooke & Woolley, 2005; Pfau et al., 2016; Matos et al., 2020), we expect that the changes induced by play in this region may also differ between the sexes. Further interrogation of the play-associated modules we identified may serve as a “window” into what mechanisms may underlie such neuroplastic changes.

We further explored potential sex-differential gene expression patterns in the amygdala in our parallel single-cell RNA-sequencing study of neonatal male and female rats. Sex-biased DEGs were enriched within inhibitory neurons in the neonatal amygdala, with relatively few seen within excitatory neurons (Figure 3c-d), consistent with what has previously been observed in the adult mouse MeA (Chen et al., 2019). GABAergic neurons within the MeA have been shown to promote various socio-sexual behaviors, including pheromone detection, reproductive behavior, and aggression (Simmons & Yahr, 2003; Pereno et al., 2011; Hong et al., 2014; Chen et. al, 2019; Lichinsky et al., 2023), while glutamatergic neurons appear to serve an antagonistic role, directing behavior away from social interactions (Johnson et al., 2021). By definition, there are strong sex differences in expression of these socio-sexual behaviors. As such, it is perhaps not surprising that such strong sex-biased gene expression patterns exist within the population of inhibitory neurons that appears to drive these sex-differential behavior patterns in adulthood. In contrast, the relative lack of sex-biased DEGs in excitatory neurons suggests that glutamatergic neurons may serve as a generalized “brake” on the system, inhibiting social behavior in a manner that does not strongly differ between the sexes.

It is also possible that further sex differences may exist within discrete subpopulations of MeA GABAergic neurons. While our scRNA-seq study was not sufficiently powered to identify sex differences within specific neuronal subtypes, our analyses revealed 11 distinct subpopulations of inhibitory neurons expressing canonical markers even at this neonatal age (Figure 3e-f), suggesting deeper sequencing studies at this timepoint may be informative. As various studies have demonstrated that aromatase-expressing cells in the MeA are critical for sex-typical social behavior (Unger et al., 2015; Yao et al., 2017), Cyp19a1+ Vgat+ cells (a population that is larger in the juvenile male MeA than that of females; Figure 5g) may represent a critical subpopulation within which sex-biased, play-induced neuroplastic changes may occur.

In our final bioinformatic experiments, we integrated our bulk and single-cell RNA-seq datasets to further investigate two representative play-associated modules of interest as case studies. Demonstrating the power of this integrative analysis, we gained useful insight on the cell-type specificity and predicted function of bM14, our representative “male play” module, and bM3, our representative “female play” module. This analysis also allowed for the identification of key hub genes in each module – *Spen* and *Klhdc8a*, respectively – which we explored further using RNAscope®. *Spen* encodes a transcriptional repressor with known roles in the estrogen signaling pathway, where it is recruited to estrogen-responsive elements upon activation of estrogen receptor alpha (Légaré et al., 2015), and as such may suggest a role for estrogen signaling in play-induced neuroplasticity in males, perhaps following local aromatization of testosterone to estradiol within the MeA inhibitory neuron population (Figure 5g). *Klhdc8a* belongs to the Kelch superfamily, representing one of the largest evolutionarily conserved gene families (Gupta & Beggs, 2014), but much less is known about its specific function. A recent study by Zhu et al. (2020) indicates that *Klhdc8a* may be involved in the MAPK signaling pathway, activation of which is believed to be a necessary component of the biochemical cascades that establish behavioral plasticity (Sweatt, 2008), including in the context of social behavior (Albert-Gascó et al., 2020). As such, this signaling cascade may be involved in mediating neuroplastic changes in the female MeA following juvenile play experience.

Together, these analyses provide novel insight into the transcriptional patterns underlying expression of social play, a fundamental adolescent behavior critical for appropriate brain development. Mirroring the qualitative and quantitative sex differences seen in expression of social play, the gene expression programs associated with play in the MeA are markedly divergent in male and female juvenile rats. We demonstrate that the inhibitory neuron population is likely central to this sex difference, as sex-biased DEGs in the neonatal amygdala are enriched within this cell type, as are play-active cells in the juvenile MeA. Through integration of multiple bioinformatic approaches, we identify key sex-biased modules and hub genes underlying sex differences in playfulness. While much remains to be discovered, our data underscore the power of an integrative, systems biology approach in exploring the underpinnings of this complex, ethologically relevant behavior.

## Acknowledgements

This work was funded by National Institutes of Health grants F31MH123025 to A.E.M., R01DA039062 to M.M.M., and S10OD026698 to the University of Maryland Center for Innovative Biomedical Resources Core Confocal Facility. We thank the University of Maryland School of Medicine Center for Innovative Biomedical Resources Core Confocal Facility (Baltimore, Maryland), including J. Mauban, for technical assistance. Sequencing was performed by the Maryland Genomics core facility within the Institute for Genome Sciences at the University of Maryland School of Medicine.

## Author Contributions

A.E.M., S.A.A., and M.M.M. conceptualized the project. A.E.M. performed the bulk RNA-seq, scRNA-seq, and Figure 6 RNAscope® experiments, and analyzed behavioral and RNAscope data. M.B. and S.A.A. analyzed all sequencing data and created relevant figures. J.V.R. conceptualized and conducted Figure 5 RNAscope® experiments and analyzed relevant data. A.E.M. wrote the original draft of the manuscript, with contributions from M.B., J.V.R., S.A.A., and M.M.M. All authors read, revised, and approved the final manuscript.

## Declaration of Interests

The authors declare no competing interests.

## Supplementary Files

Supplementary Figures S1 through S5

Supplementary Tables S1 through S9

## Notes

### Competing Interest Statement

The authors have declared no competing interest.

