## Supplementary Figures S1-S9 and Table Legends for "The transcriptome of playfulness is sex-biased in the juvenile rat medial amygdala: a role for inhibitory neurons"

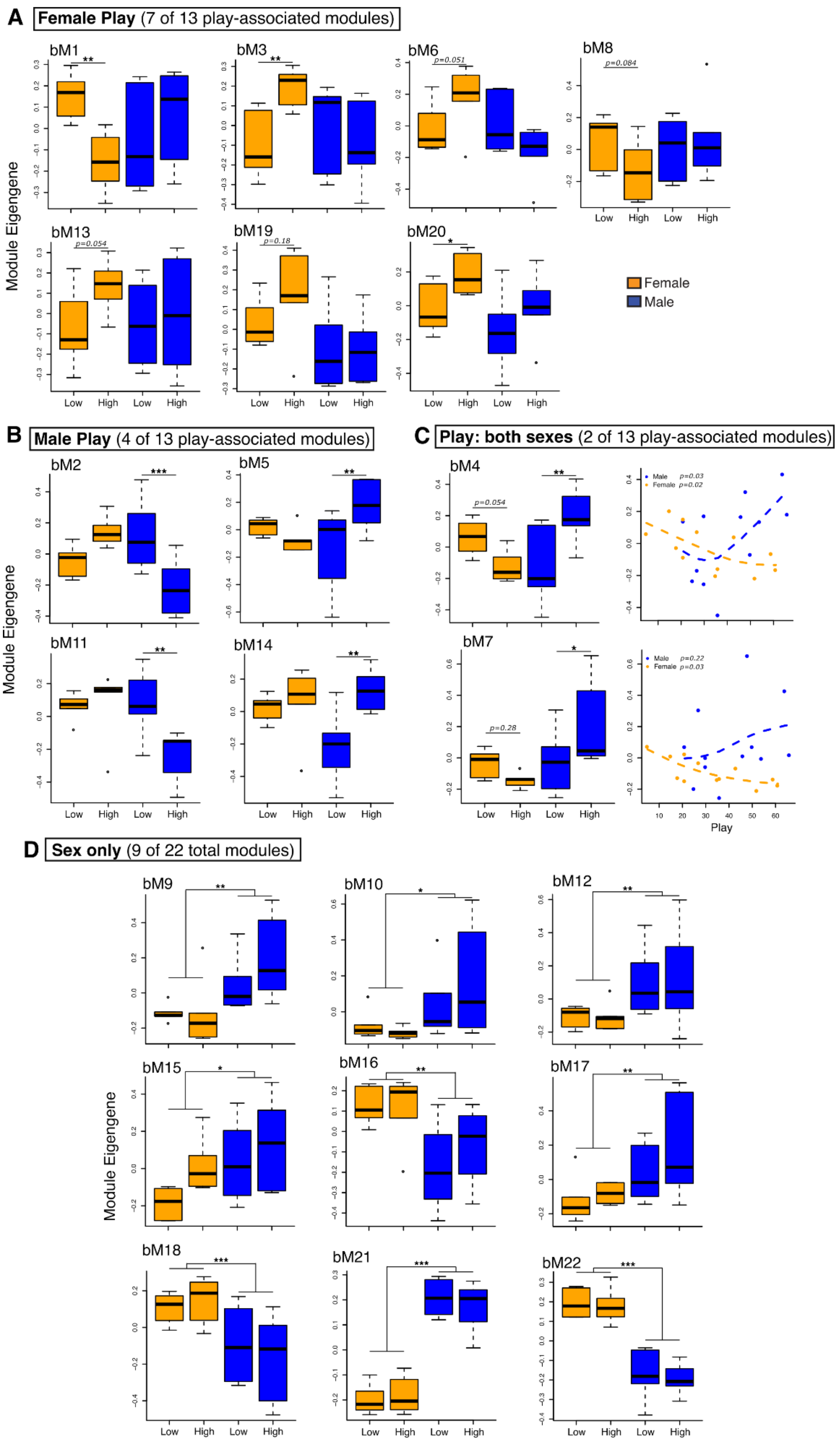
A Female Play (7 of 13 play-associated modules)
bM1
bM3
bM6
bM8
bM13
bM19
bM20
p=0.051
p=0.084
p=0.054
p=0.18
Female
Male
Module Eigengene
Low
High
B Male Play (4 of 13 play-associated modules)
bM2
bM5
bM11
bM14
C Play: both sexes (2 of 13 play-associated modules)
bM4
bM7
p=0.054
p=0.28
Male p=0.03
Female p=0.02
Male p=0.22
Female p=0.03
Play
D Sex only (9 of 22 total modules)
bM9
bM10
bM12
bM15
bM16
bM17
bM18
bM21
bM22

**Figure S1. Association of all 22 bulk RNA-seq modules with sex and play.** Module eigengene expression across groups for modules deemed to be associated with Female Play (***a***), Male Play (***b***), Play: both sexes (***c***), or Sex only (***d***). The association of the module eigengene with quantitative play score is also shown for the two modules associated with play in both sexes (***c***, right panel). “Low” = low-playing; “High” = “high-playing”. *$p < 0.05$, ** $p < 0.01$, ***$p < 0.001$, $n = 6$ per group.

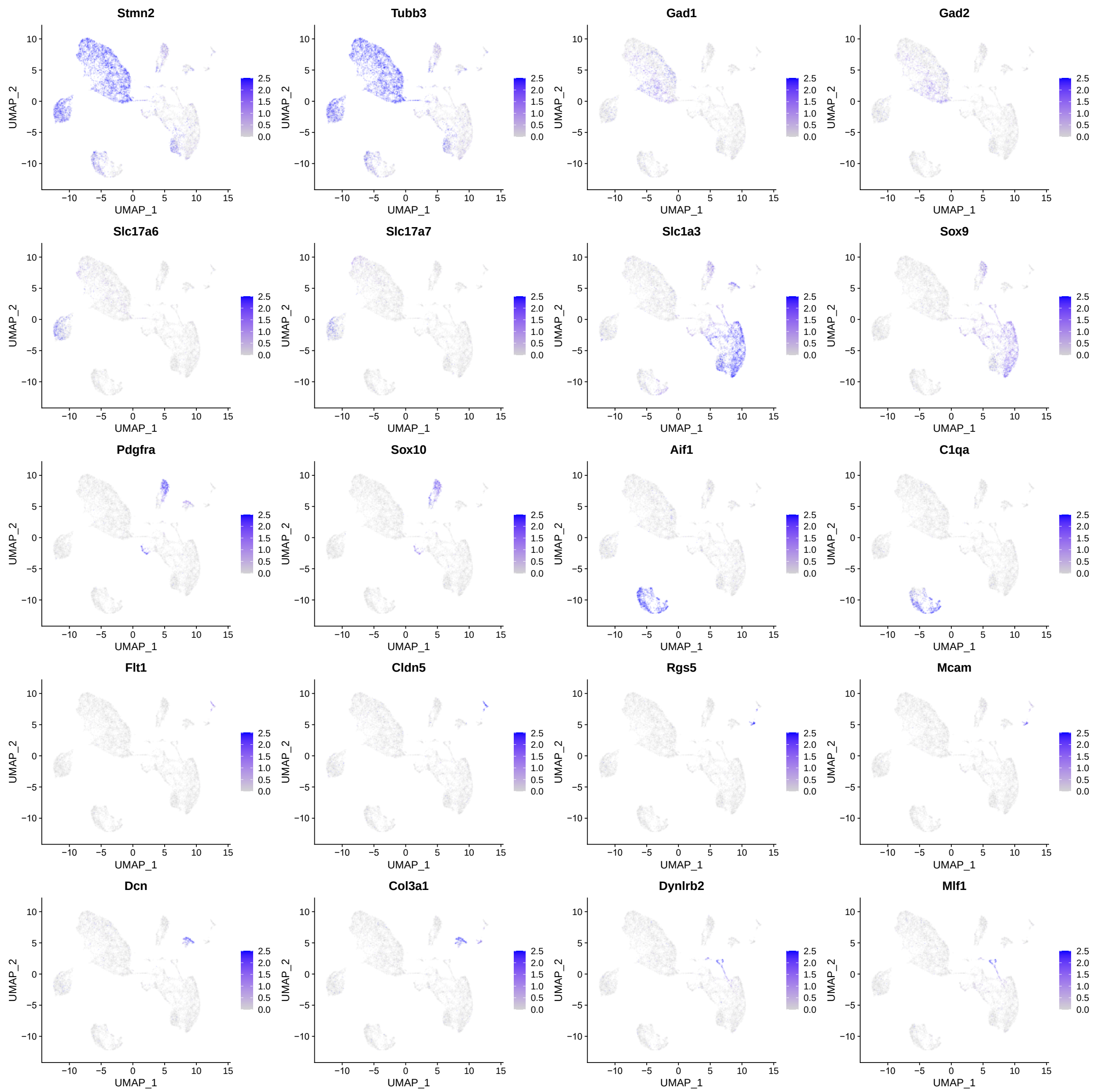


**Figure S2. Expression patterns of cell type marker genes from scRNA-seq experiment.**

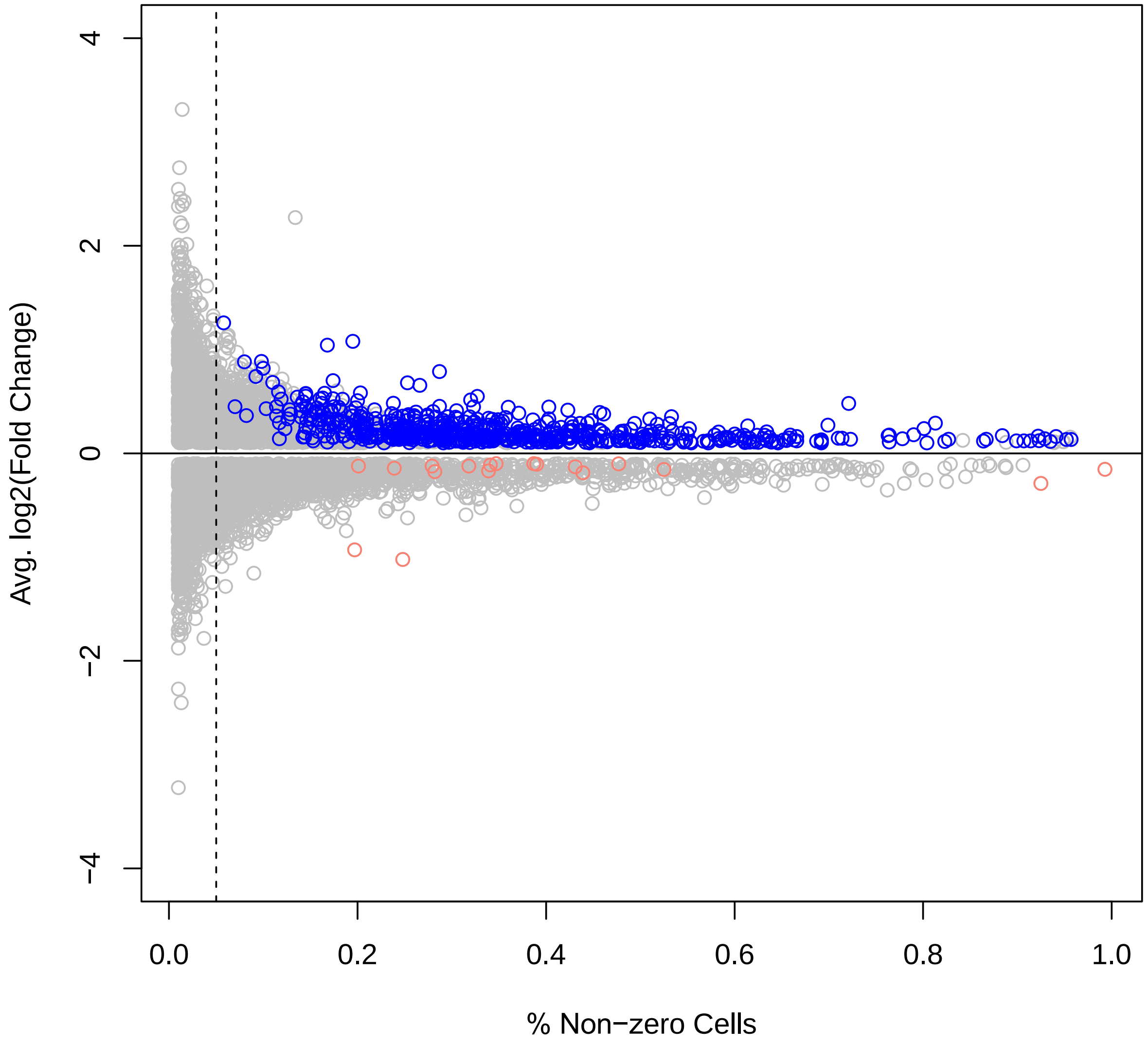


**Figure S3. Quality control of inhibitory neuron differential gene expression statistics.** The variance and distribution of fold change estimates is informative about the reliability of differential gene expression statistics. We plotted the log2(fold change) estimates in inhibitory neurons from males vs. females (y-axis) as a function of expression level (% of cells with at least one sequencing read; x-axis). The variance in fold changes was much wider for genes detected in <5% of cells (dotted line), so these were filtered out from downstream analyses. Importantly, although we detected more male-biased (blue) than female-biased (pink) differentially expressed genes (FDR < 0.01), the background distribution of non-significant fold changes was not skewed, with similar numbers of genes displaying non-significant (random) variation above and below zero. Note: This plot excludes X and Y chromosome genes because their distributions are expected to be biased between sexes.

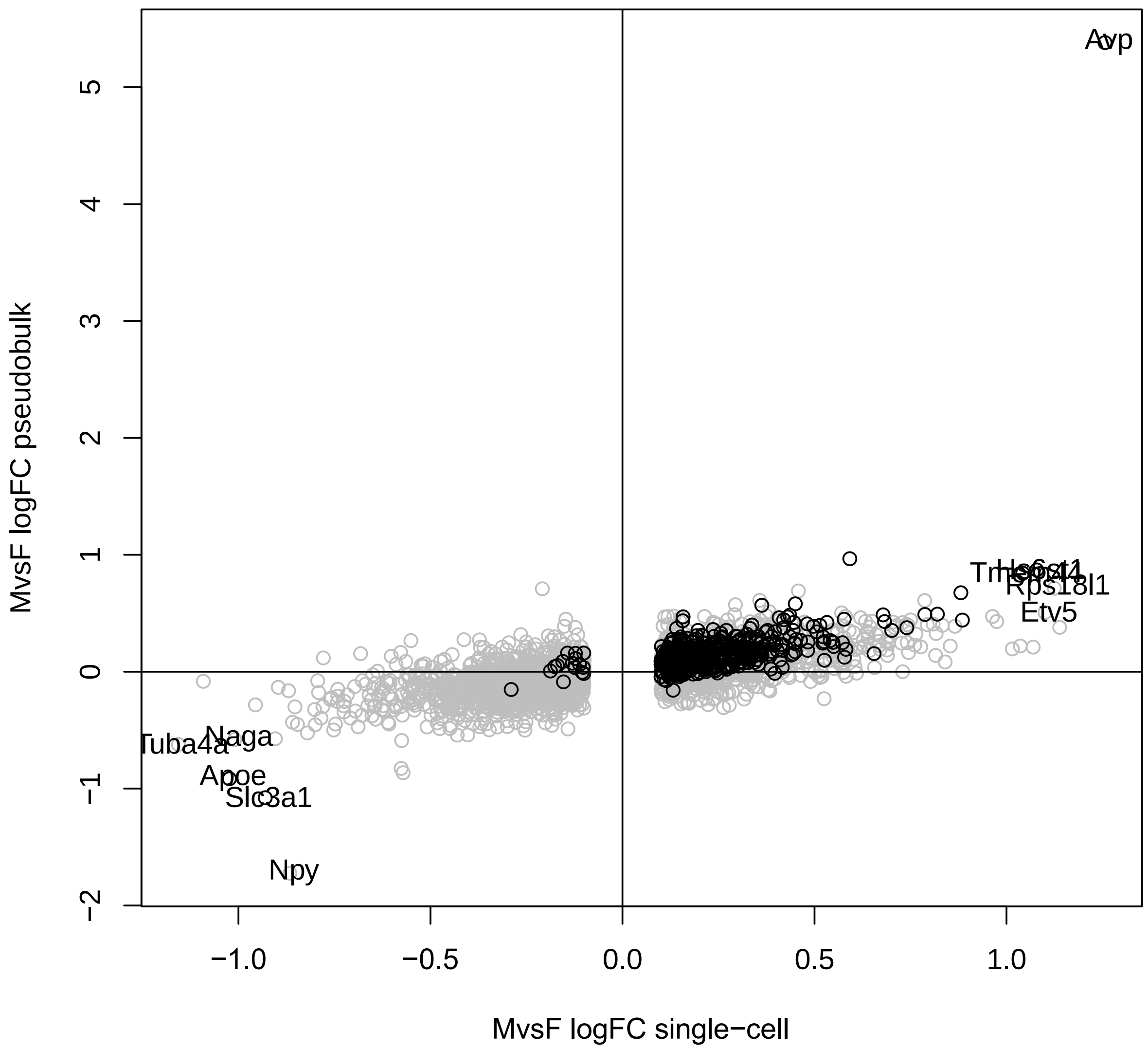


**Figure S4. Estimates of sex-biased gene expression in inhibitory neurons are robust to analytical approach.** In our primary analysis, we assessed sex differences in gene expression based on read counts in individual cells. As an alternative approach, we used the summed read counts from all the cells of each cell type within each independent biosample (cells pooled from three male or female animals). The latter approach is underpowered to assess statistical significance in this dataset but is more robust to batch effects. The log2(fold change) from the first method is shown on the x-axis, and the log2(fold change) of the second method is shown on the y-axis. Genes with significant p-values in our primary analysis are indicated by black-colored points. Overall, we found that the fold-change estimates from these two approaches were strongly correlated (r = 0.65), and the top DEGs (labeled genes) were reliably detected by both approaches.

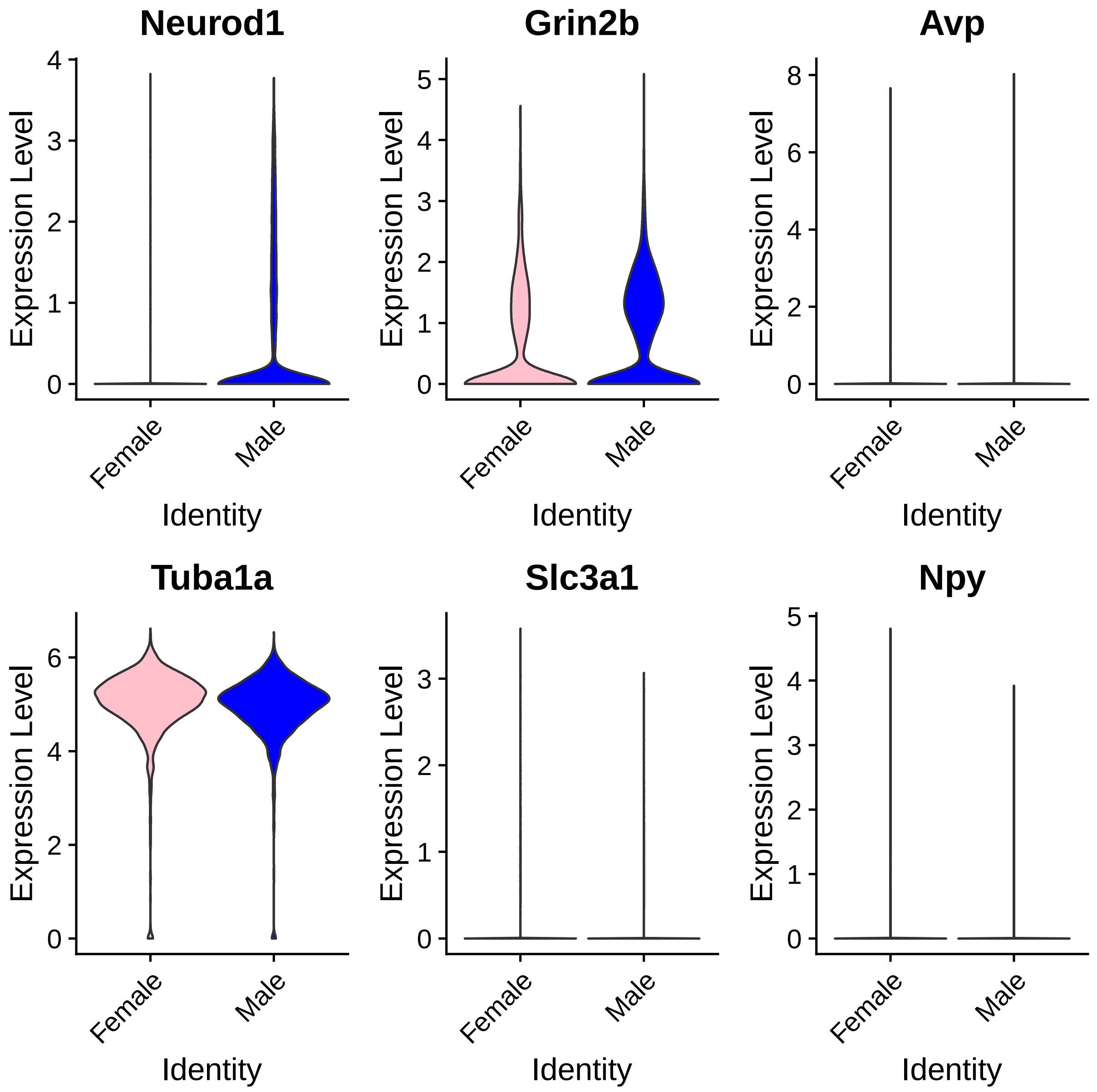


**Figure S5. Expression patterns of selected sex-biased genes in inhibitory neurons.** Neurod1 and Grin2b were among the top male-biased genes by p-value, and *Tuba1a* and *Slc3a1* were among the top female-biased genes by p-value. *Avp* and *Npy* are expressed in only a small proportion of inhibitory neurons but had amongst the largest reported fold differences in males vs. females.

**Supplementary Tables**

| Number | Description | Relates to |
|---|---|---|
| Table S1 | Bulk RNA-seq WGCNA module assignment by gene | Figure 2 |
| Table S2 | Characterization of each bulk RNA-seq WGCNA module | Figure 2 |
| Table S3 | scRNA-seq major cell type markers | Figure 3 |
| Table S4 | scRNA-seq sex-biased DEGs – by single-cell | Figure 3 |
| Table S5 | scRNA-seq sex-biased DEGs – by pseudobulk | Figure 3 |
| Table S6 | scRNA-seq inhibitory neuron subcluster markers | Figure 3 |
| Table S7 | scRNA-seq WGCNA module assignment by gene | Figure 4 |
| Table S8 | Overlap of scRNA-seq & bulk RNA-seq WGCNA modules | Figure 4 |
| Table S9 | Association of each scRNA-seq module with sex and play | Figure 4 |
